# Aberrant prefrontal activity and arousal level correlate with action initiation and response vigor

**DOI:** 10.1101/2024.09.04.611286

**Authors:** Frederike J. Klein, Dmitrii Vasilev, Ryo Iwai, Masataka Watanabe, Nelson K. Totah

## Abstract

Orchestrating learned Stimulus-Response (S-R) mappings has been suggested as one of the central functions of the prefrontal cortex (PFC). While S-R selective activity has been demonstrated, it remains unclear whether the strength of such activity is related to the vigor of the subsequent response. Here, we trained male rats to perform a Go/NoGo response task while head-fixed on a treadmill. This allowed us to record PFC (cingulate, area 24) single unit spiking, as well as running speed as a proxy for response vigor. We show that aberrant activation of the “wrong” S-R mapping is correlated with initiation of the incorrect response. The vigor of the incorrect response was directly related to the strength of the aberrant stimulus-evoked activity. A similar relationship was observed for pre-trial arousal state and response vigor. Our findings confirm the long-standing concept, established in psychology and cognitive neuroscience, that S-R mappings are directly related to response vigor. Moreover, we provide evidence for the often suggested but rarely tested relationship between arousal and response vigor and a potential underlying neuronal mechanism involving neuromodulation of S-R mapping activity.

**Significance statement:** The concept of stimulus-response (S-R) mappings is fundamental in psychology and has been widely documented as a key function of the prefrontal cortex. Here, the authors directly link prefrontal single neuron mapping-selective activity to the vigor of responses. Moreover, they link a physiological measure of arousal to response vigor and suggest that neuromodulatory systems invigorate responses by potentially modulating PFC S-R mapping-selective activity.

## Introduction

The brain encounters a myriad of stimuli in any given moment. While not all these stimuli are used to control behavior, some of them are associated with specific responses. Think, for example, of a traffic light turning red and the potentially catastrophic consequences of failing to select the correct response: stopping. Stimulus – Response (S-R) mappings, like stopping at a red light, are often learned associations. Miller & Cohen (2001) suggested that a primary function of the prefrontal cortex (PFC) is to represent learned S-R mappings and thus, select and initiate the appropriate stimulus-guided response. According to this theory, specific groups of PFC neurons should be activated after presentation of a particular stimulus, S1, but only if the subject commits the correct stimulus-associated response (R1) and not when the subject makes the incorrect response (R2). Hence, these neurons represent the S1-R1 mapping.

In line with this hypothesis, when a stimulus-response mapping is learned, subsets of PFC neurons in non-human primates become selectively responsive to specific stimuli (Niki, 1974; Watanabe, 1986; Yamatani et al., 1990; Sakagami and Niki, 1994a; Bichot et al., 1996; Miller et al., 1996; Sakagami and Tsutsui, 1999). Moreover, the stimulus-evoked activation of those neurons that acquire stimulus-selectivity occurs only when the subject commits the response associated with that stimulus. Similar neuronal correlates of S-R mappings have been observed in the PFC of humans (Woolgar et al., 2011a, 2011b), rodents (Schoenbaum and Eichenbaum, 1995; Peters et al., 2005; Reinert et al., 2021; Wal et al., 2021), and the nidopallium caudolaterale of corvids (Veit and Nieder, 2013; Moll and Nieder, 2015; Veit et al., 2015).

Classically, the activity of S-R mapping selective PFC neurons has been described for trials in which the subject performed the correct response. Single neuron examples from error trials show that, in these trials, the “wrong” PFC neurons were aberrantly activated (Veit and Nieder, 2013; Moll and Nieder, 2015; Veit et al., 2015; Chen et al., 2017; Schmitt et al., 2017). In other words, PFC neurons tuned to the S2-R2 mapping aberrantly responded after presentation of the other stimulus, S1, and the subject subsequently committed an error (i.e., the response, R2). We hypothesize that the strength of this aberrant activity and the “strength” or vigor of R2 is correlated.

Notably, while psychologists and cognitive neuroscientists (e.g., Miller & Cohen in 2001) have predicted that the neural representation of an S-R mapping should relate to the vigor of the response, this has neither been tested nor demonstrated. Most of these classic S-R mapping PFC studies (Watanabe, 1986; Sakagami and Niki, 1994a, 1994b; Sakagami and Tsutsui, 1999; Veit and Nieder, 2013; Moll and Nieder, 2015; Veit et al., 2015; Chen et al., 2017; Schmitt et al., 2017) have made use of behavioral tasks in which the stimulus presentation and the subject’s response are separated by a delay period. The mapping selective activity is observed during the delay period and is no longer apparent at the time of response initiation. This has made it impossible to relate the mapping selective activity to action initiation or the vigor of the subsequent response.

Here, we investigate the relationship between the vigor or strength of a stimulus-guided response and the preceding mapping selective activity in the rat PFC. We find that the trial-specific strength of aberrant activity is directly correlated to incorrectly initiated running speed. We further demonstrate that a similar relationship can be observed for pre-stimulus pupil size and the running speed. These findings suggest that the extent to which a specific S-R mapping is activated in PFC can be directly related to the strength of response that is initiated, and that response vigor in this context is modulated by pre-trial arousal state.

## Material and Methods

### Subjects

Male Lister-Hooded rats (140-190 g body weight) were used for single unit recordings (N=3) and pupillometry (N=37). The rats were supplied by Charles River Laboratories (Germany) and were housed in pairs for 7 days prior to implantation of the head-fixation implant. After implantation, rats were single housed on a reversed light-dark (07:00 lights off, 19:00 lights on) cycle. Training and experiments were performed during the rats’ active phase. All procedures were carried out after approval by local authorities and in compliance with the German Law for the Protection of Animals in experimental research (Tierschutzversuchstierverordnung) and the European Community Guidelines for the Care and Use of Laboratory Animals (EU Directive 2010/63/EU).

### Surgery

The animal was anesthetized with isoflurane (∼1.0 – 2.0%). Heart rate was monitored throughout surgery. Buprenorphine (0.06 m/kg, s.c.), meloxicam (2.0 mg/kg, s.c.), and enrofloxacin (10.0 mg/kg, s.c.) were administered. An incision was made once the rat was no longer responsive to paw pinch. Skin and connective tissue were removed to expose the skull from the frontal bone to the neck muscle and from left to right temporal muscles. The wound margin was cauterized. The exposed bone was wiped dry and cleaned with 5% hydrogen peroxide. The bone surface was then scratched with a bone curette in a grid pattern to facilitate adhesion of the adhesive for the UV light polymerizing cement used to affix the implant to the skull. Two component UV-curing adhesive (OptiBond, Kerr) was applied to the skull and UV cured for 30 sec at full intensity (Superlite 1300, M+W Dental). A custom-made head fixation implant was attached to the skull using UV-curing cement (Tetric EvoFlow, Ivoclar). The dental cement was bonded to the adhesive by UV curing for 60 sec at full intensity. In rats that were not implanted with a multi-electrode silcon probe, the chamber was filled with 2-component dental cement (Paladur, Kulzer). In cases where multi-electrode probes were to be implanted, the skull was covered in biocompatible silicone elastomer (KwikCast, WPI) and the implant was closed using a lid and screws. In all rats, the skin was glued to the sides of the implant using tissue glue (Histoacryl, B. Braun).

Rats that were implanted with a multi-electrode silicon probe were first trained in the behavioral task. The rats then underwent a second surgery, in which the lid of the implant and the silicone elastomer was removed. The probe (one rat, A2×32, Neuronexus; two rats, H9×64, Cambridge Neurotech) was implanted into the PFC (target coordinates from bregma: AP: 2.7 mm; ML: 0.8 mm; depth: 3.2 – 4.4 mm, varying across rats) through a craniotomy. Additionally, a second craniotomy was made over the cerebellum. A reference electrode (99.9% pure silver wire) was inserted through this posterior craniotomy. The craniotomy was filled with viscous, electrically-conductive agar. The open space on the skull was filled with 2-component dental cement (Paladur, Kulzer).

Rats recovered for 5 days after surgery Buprenorphine (0.06 m/kg, s.c.) was administered every 12 hours for 3 days in some rats and other rats were injected with meloxicam (2.0 mg/kg, s.c.) every 24 hours for 3 days. Rehydrating and easily consumable food was provided (DietGel Recovery, ClearH2O).

### Handling and water restriction

For five days prior to surgery, the rats were handled twice a day, once in the morning and in the evening. Each session lasted at least five minutes. After five days of post-surgical recovery, access to water was restricted. During training and experiments, the rats were given 8-12 mL total water per day. Most of the water was consumed as reward during the behavioral task. The remainder of the total water volume was supplied to the rats in the cage after training. The total volume of water available daily was restricted to this level for between 5 and 14 days, while rats learned and performed stimulus discrimination experiments. After an epoch of restricted water availability, rats were provided ad libitum access to water for 24 hours.

### Head-fixation and behavioral apparatus

The rat was head-fixed on a cylindrical, non-motorized fibreglass treadmill that rotated forward or backward freely on low-friction ball bearings. The treadmill and head-fixation apparatus were inside a large Faraday cage (approximately 2 m x 2 m x 2 m) with sound proofing material. A TTL pulse-controlled pump was used to deliver 10% sucrose water via a reward port that was placed at the mouth of the rat. A computer screen (behind glass with electromagnetic shielding designed to not cause a Moire effect) in front of the rat was used to display visual stimuli covering the entire visual field of the rat. Treadmill angular position was recorded via an analog signal output from a rotary encoder (MA3-A10-125-B, US Digital) attached to the rotational axis of the cylindrical treadmill. The signal output varied between 0 V and +5 V, which mapped linearly to the rotational angle of the treadmill. The signal was sampled at 32 kHz, digitized (Neuralynx signal acquisition system), and velocity was calculated offline (in MATLAB). Video for pupillometry was recorded from the right eye at 45 fps under near-infrared illumination (M850L3, Thor Labs LED, with COP4-B, Thor Labs collimation optics). Frames were recorded with a near-infrared camera (G-046B, Allied Vision) and a variable zoom lens, fixed 3.3x zoom lens, and 0.25x zoom lens attached in-line (1-60135, 1-62831, 6044, Polytec). The camera provided a TTL pulse with each video frame, which was recorded by the Neuralynx signal acquisition system at 32 kHz.

### Habituation to head-fixation and behavioral task training

Habituation to head-fixation consisted of a single 20 minute session. After habituation, rats were trained to commit an instrumental response for reward. Approximately 5 uL of reward solution (10% sucrose in water) was delivered for small “shaking” or body movements on the treadmill. The threshold for triggered reward was gradually increased to train the rat to make larger body movements and eventually steps. Threshold crossings were marked with a bridging stimulus (0.1 sec duration, 500 Hz auditory tone) to aid in learning the link between movement and reward. Eventually, rats would continuously walk and receive reward. This stage required from 3 to 10 sessions (one per day). Once an animal was running and licking simultaneously (which yielded approximately 7 mL of reward solution in a session lasting 20-30 minutes), we trained the rat to make instrumental (Go) responses contingent upon the presentation of a visual stimulus.

Initially, we presented a 15 sec duration visual stimulus. The stimulus was a full field, black and white drifting grating (2.4 cycles/sec, 0.005 cycles/pixel spatial frequency, 75 deg orientation). Rats were trained to respond to the stimulus by continuously delivering reward for running during stimulus presentation. Reward delivery was triggered by crossing a threshold (a.u.) that was the same for all rats and all sessions and set at a level that was associated with bilateral locomotion. The stimulus was followed by an inter-trial interval (ITI). The ITI duration was drawn randomly from a distribution ranging from 2 to 3 sec (0.05 sec bins size).

After 2 sessions, the rats were trained to not respond prior to stimulus onset. The ITI was reduced to 1 to 2 sec, and any running that crossed a velocity threshold (manually set to capture running, same for all rats and sessions) resulted in a 0.5 sec time-out from the task and a resetting of the ITI. After one or two sessions, the rats started to suppress running during the ITI. Once this was achieved, we reduced stimulus duration in small steps (10 sec, 5 sec, 2.5 sec) over a few sessions. When stimulus duration is 2.5 sec, rats exhibit a vigorous and low-latency response upon stimulus onset. At this point in training, we reduce the ITI to 0.5 to 1 sec and, after a few sessions, we reduce stimulus duration to 1.5 sec (i.e., a speeded reaction time task). Rats were given 600 trials per session. This typically yielded approximately 6 mL of sucrose solution during the task. Behavior was considered stable when omission rate was below 10%.

The Go/NoGo paradigm was introduced with the addition of a NoGo stimulus. The NoGo stimulus was at least 70 degrees different from the Go stimulus. Go and NoGo stimulus trials were delivered in pseudo-random order and in equal proportion. At most, two trials of the same stimulus type could occur consecutively. A Go response required crossing a distance threshold, which roughly corresponded to taking one step. A response offset window (0 to 0.75 sec after stimulus onset) was introduced to compensate for the pre-potent drive to respond. During this period, running did not count towards the distance threshold. This allowed low latency movements but forced the rat to appraise the stimulus and make a decision. After the offset window, crossing the distance threshold caused the stimulus to disappear. Hits were rewarded (three 7 uL pulses). Responses to the NoGo stimulus led to an auditory error signal (0.5 sec duration, brown noise, 60 dB) and a time-out of 6 seconds prior to the next ITI. Training was complete when performance was above 85% and omission rate was less than 10%.

### Neurophysiological recordings and spike sorting

Wideband (0.1 Hz to 10 kHz) signals were recorded at 32 kHz (Digital Lynx SX, Neuralynx). Automatic spike sorting was performed using KiloSort 4.0 (Pachitariu et al., 2024). Afterwards, the outputs were manually curated using standard criteria (i.e., stable firing rate, waveform similarity, auto- and cross-correlograms).

### Single unit spike count analysis

A Go stimulus-preferring unit responded to the Go stimulus and not to the NoGo stimulus on “pure” Correct Rejection trials, which had little-to-no running. Spike count peri-stimulus time histograms (0.1 sec bin size, -0.5 sec to 1.5 sec window around stimulus onset) were z-scored to the trial-averaged spike counts in the 0.5 sec before stimulus onset for each unit. The unit was considered responsive to the stimulus, if the z-score was greater than 2 for three consecutive bins in the post-stimulus window (0 to 1.5 sec).

We quantified the peak activity of units as the maximal z-scored spike count in the peri-stimulus time histogram. The Hit/Omission index was calculated as:

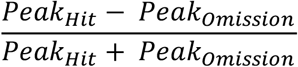

### Pupil analysis

Pupillometry was implemented using a custom computer vision algorithm built with the openCV package in Python 3.7. A detailed description of the method is in our prior work (Vasilev et al., 2023).

### Experimental Design and Statistical Analysis

This study used 40 male Lister-Hooded rats (3 for single unit recordings and 37 for pupillometry). Bayesian statistics (JASP software) were used to assess evidence in favor of the null hypothesis and in favor of the alternative hypothesis (Keysers et al., 2020). We report BF10 which reports the evidence in favor of the alternative hypothesis over evidence favoring the null hypothesis. All analyses used a one-way ANOVA. The single unit activity data in Figure 3A were analyzed with a repeated-measures ANOVA because the same single units were compared across 3 conditions. The trialwise velocity data in Figure 4C were analyzed with an independent samples ANOVA because there were different trial numbers within the 3 conditions (small, medium, and large pupil size) that were being compared. This is due to pupil size freely varying across sessions and rats.

## Results

We recorded activity in the PFC of three male rats during a visually cued Go/NoGo task (**Figure 1**). A total of 629 single units were recorded. Approximately 9% (58 out of 629 units) selectively responded to the Go stimulus in correctly performed trials. This means that they had a clear stimulus-evoked response in Hit trials, while they were not activated in Correct Rejection (CR) trials. Our analyses were focused on these 58 Go stimulus-preferring units, as they are responsive to one S-R mapping, Go stimulus (S1) – running (R1), and not responsive to the other mapping, NoGo stimulus (S2) – immobility (R2).

**Figure 1.**
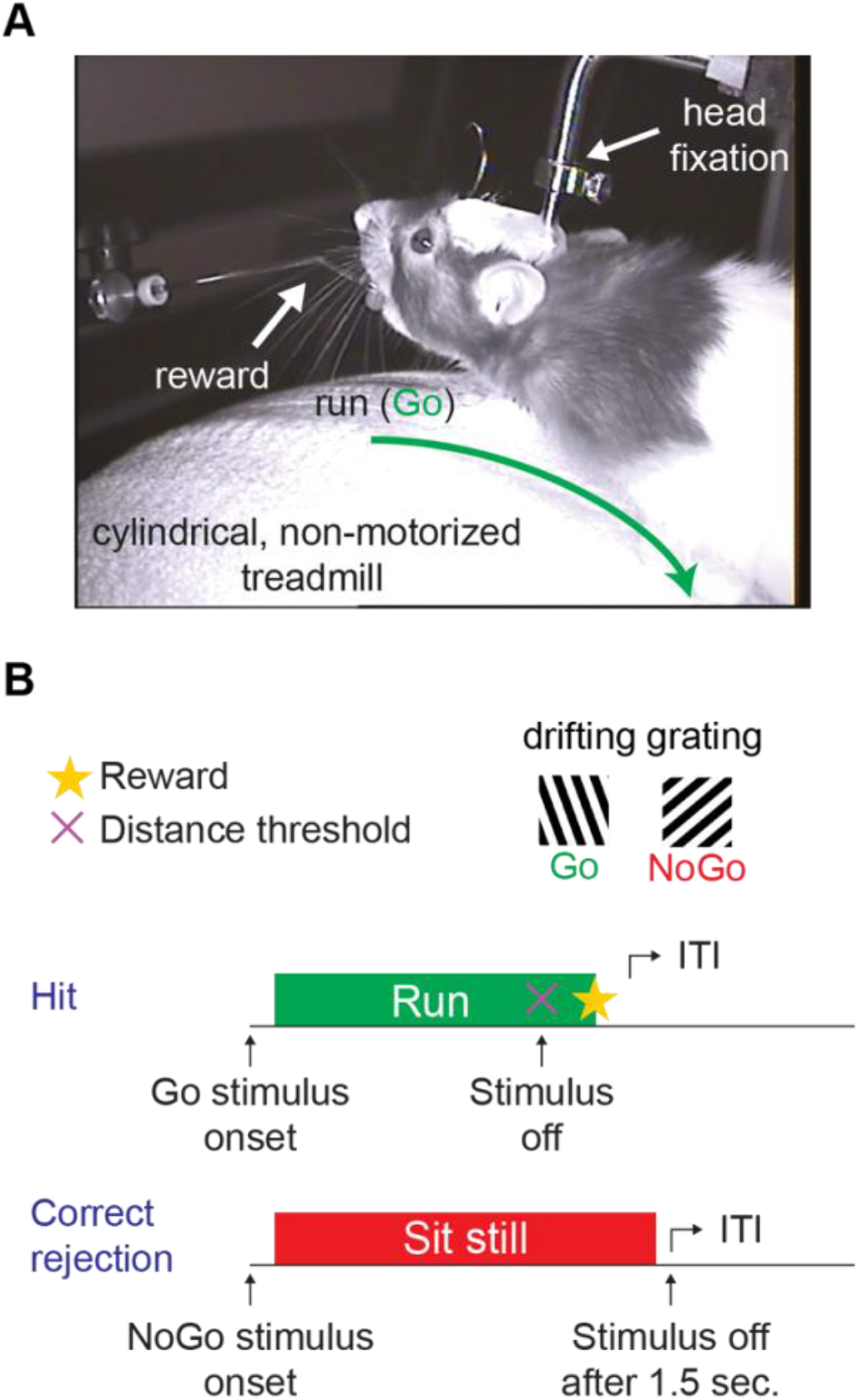
Set-up and task design. A Illustration of the set-up. Rats were head-fixed on a cylindrical, non-motorized treadmill. A reward spout was positioned in front of the rat’s mouth through which 10% sucrose solution was delivered on Hit trials. **B** A schematic showing trial progression for Hit and Correct Rejection trials in the Go/NoGo task. The Go and NoGo stimuli consisted of a drifting grating with different orientation. Rats were required to sit immobile for at least 0.5 sec prior to stimulus onset. In a Hit trial, the rat started running upon presentation of the Go stimulus until a distance threshold was crossed and reward was delivered. On trials in which the NoGo stimulus was presented, the rat was supposed to sit still until the stimulus was turned off for a succesful CR response.

### Activity in CR trials is correlated with incorrect action initiation

Despite Go-preferring units being primarily responsive in Hit trials relative to CR trials, the spike rasters made it apparent that individual Go stimulus-preferring units were also active in a subset of CR trials (**Figure 2A**). This aberrant activity appeared in trials in which the non-preferred (i.e., NoGo) stimulus was presented and in which the distance threshold for a False Alarm was not crossed. We examined the response trajectory on trials with these aberrant activations. To do so, we averaged the post-stimulus velocity across these trials and compared to trials without activation (0 spikes during the 1 sec window after stimulus onset). This analysis revealed that when the unit aberrantly activated, running was incorrectly initiated, but aborted before the rat crossed the distance threshold thus avoiding a False Alarm (**Figure 2B**). This finding indicates that activation of the aberrant S-R mapping is associated with the initiation of an incorrect response.

**Figure 2.**
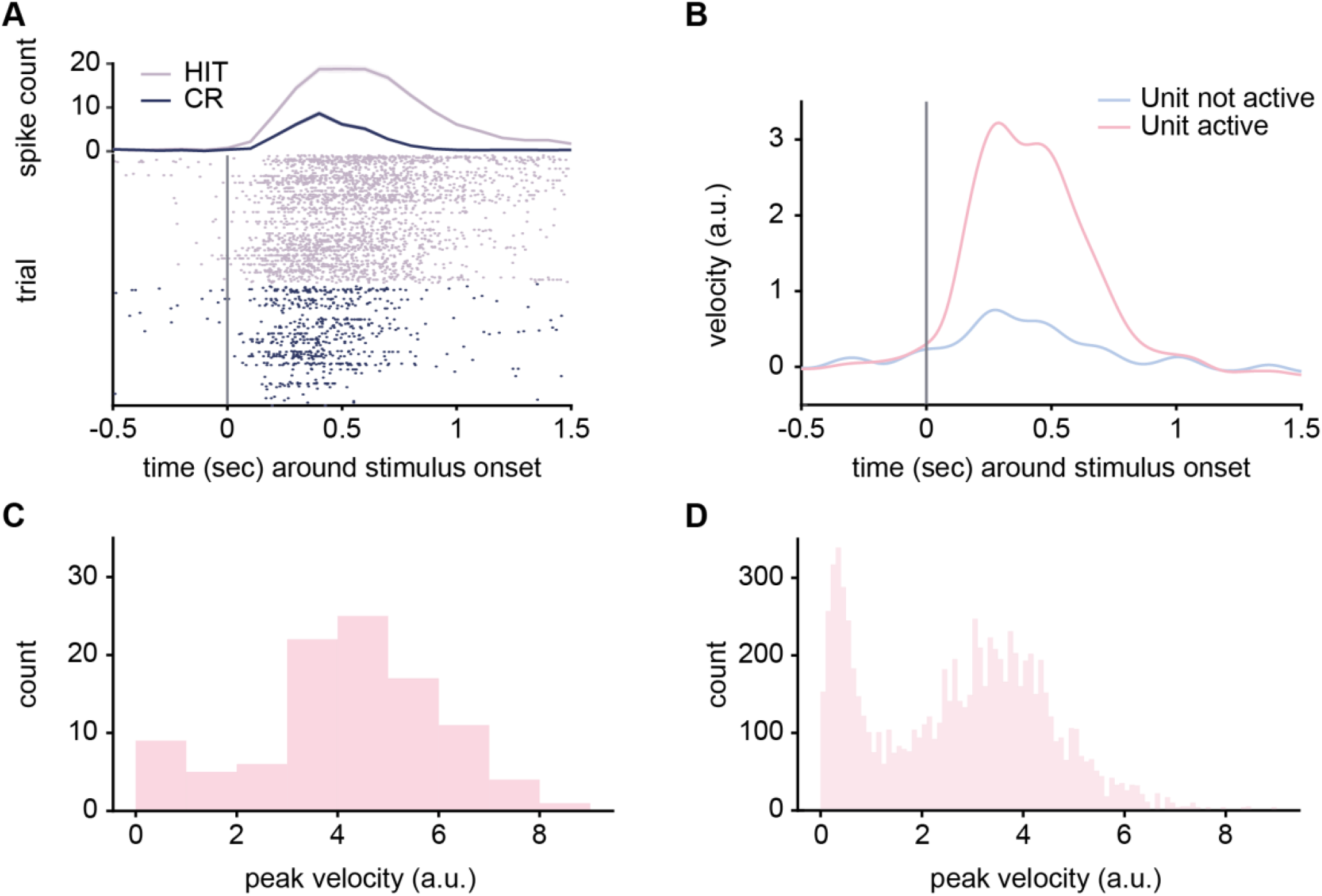
Activity of a Go stimulus-preferring unit and running behavior on CR trials. A Example single unit spike raster plot (bottom) and peri-stimulus time histogram (top) for HIT and CR trials. The grey line marks the time of stimulus onset. **B** Velocity of the rat averaged across CR trials in which the example unit, shown in panel A, was active (pink) or not active (light blue). Trials in which the unit was active showed a brief period of running after NoGo stimulus presentation. **C** A histogram of maximum velocity for each CR trial in which the example unit was active. This revealed a wide variability in peak velocities. **D** A histogram showing peak velocity (same as in panel C), but for all 58 Go stimulus-preferring units.

### Strength of aberrant activity is correlated with response vigor

In the session in which this example unit was recorded, we observed that peak velocity varied across CR trials (**Figure 2C**). Since CR trials with aberrant activity were present across the population of 58 Go stimulus-preferring units, we assessed whether peak velocity varied at the level of the neuronal population. Similar to the example unit shown in **Figure 2A-C**, trials in which the individual units were aberrantly active were accompanied by incorrect initiation of motion with a variance in peak velocity (**Figure 2D**) before the animals stopped for an eventual CR. Since both the stimulus-evoked spike rate and peak velocity varied, we investigated whether the strength of activity was correlated with running speed (i.e., response vigor) on a trial-by-trial basis.

We investigated the relationship between the variability in neuronal activity and peak velocity on CR trials by splitting the trials into low, medium, and high peak velocity groups. As velocity increased, the NoGo stimulus-evoked activity increased (**Figure 3A**). A Bayesian repeated-measures ANOVA supported the alternative hypothesis of an activity difference depending on velocity (BF_10_ = 7816.95). Bayesian post-hoc t-tests supported an obvious difference between low velocity compared to medium and high velocity trials (BF_10_ = 9689.83 and BF_10_ = 212.78, respectively). The Bayesian analysis suggested that there is moderate evidence in favor of increased activity in high velocity trials compared to medium velocity trials (BF_10_ = 2.85). At the level of individual units (**Figure 3B-D**), as response vigor increased, the activation gradually approached the level of activity on Hit trials. This effect was most prominent for units with higher Hit trial spike count. This suggests that the level to which aberrant S-R mapping activity is present in PFC is directly related to the vigor with which the incorrect response is initiated in these trials.

**Figure 3.**
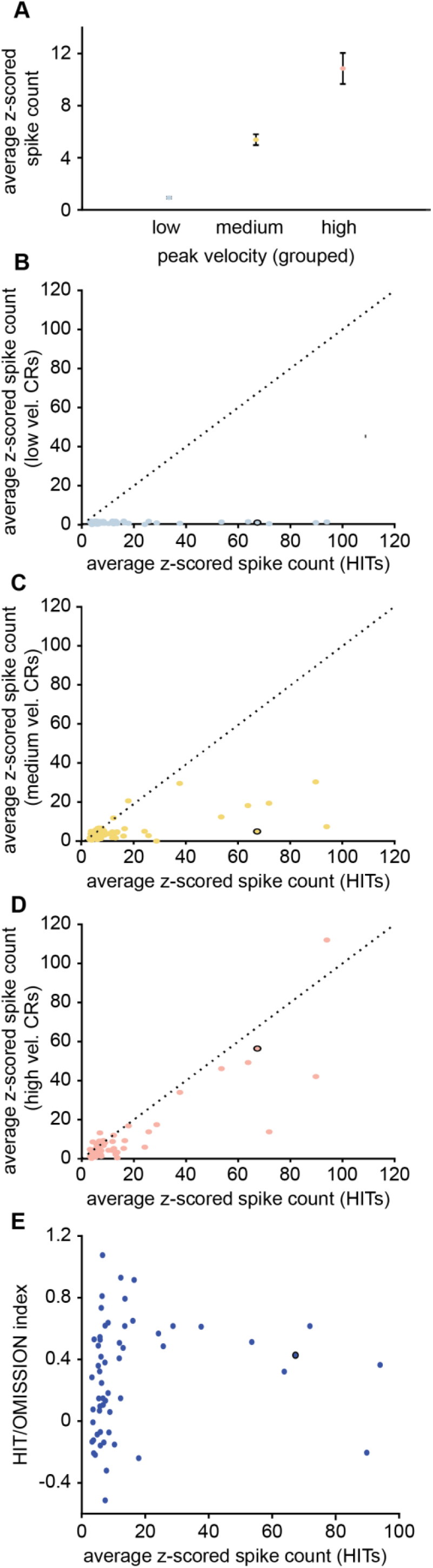
Relationship between peak velocity and stimulus-evoked spike counts. A Trials were grouped into sets with high, medium, and low peak velocity. Average z-scored spike count increased with peak velocity. The low peak velocity group was different from both the medium (BF_10_ = 9689.83) and high (BF_10_ = 212.78) peak velocity groups. Evidence for a difference between medium and high peak velocity groups was only moderate (BF_10_ = 2.85). **B** Z-scored spike count for HIT trials (x-axis) and low peak velocity CR trials (y-axis) for each of the 58 Go stimulus-preferring units. The example unit from **Figure 2A-C** is marked with a black circle. **C** Same as B, but for medium peak velocity CR trials. **D** Same as B, but for high peak velocity CR trials. **E** Hit/Omission index values for all Go stimulus-preferring units plotted against z-scored Hit trial spike counts. Index values <= 0 indicate selectivity for the Go stimulus rather than selectivity for the S-R mapping.

While most of the Go stimulus-preferring units in our study responded on CR trials, when incorrect running was initiated, some units exhibited little-to-no increase in spike count. These units tended to fire at a lower rate on Hit trials. Thus, they might have only a slight preference for the Go stimulus, and given their low activity level scaling with response velocity could be difficult to observe. On the other hand, it is possible that these units are not representing the S-R mapping but rather developed a preference for the Go stimulus itself. In this case, we would not expect any scaling and instead expect the units to remain unresponsive to the NoGo stimulus. To test this, we computed a Hit/Omission index (see Methods). If a unit is selective for the Go stimulus itself rather than the S-R mapping, then it should also be active in Omission trials since the same stimulus is presented. If a unit is not encoding the S-R mapping, then the index will be close to 0, or negative. For S-R mapping selective units, we expect a higher index value since these units should not be active in Omission trials (when no response is initiated). We indeed observed a group of units with negative Hit/Omission index values, which were almost exclusively those with low Hit trial spike counts (**Figure 3E**). Therefore, we can assume that a subset of the Go stimulus-preferring units are purely selective for the Go stimulus itself rather than the mapping of this stimulus to running. This explains the lack of response scaling with running speed in CR trials for those units.

### Pre-stimulus arousal is related to peak velocity

One factor that has been shown to influence response speed or reaction time and thus response vigor is arousal. We used a dataset including pupillometry data from 37 rats performing the same Go/NoGo task. Pupil size is a proxy for the arousal state (Bradshaw, 1967). We focused on pupil size in a 0.5 sec window prior to stimulus presentation in CR trials (see **Figure 4A** for an example). Pupil size was then averaged across this time window. This revealed variable pupil sizes across the duration of a single recording session (**Figure 4B**). We tested whether pre-stimulus pupil size was correlated with the vigor of incorrectly initiated responses by splitting the trials into three groups based on pupil size (small, medium, and large pupil). Peak velocity for small pupil trials was significantly slower than for the other two pupil size conditions (**Figure 4C**). A Bayesian independent samples ANOVA supported an effect of pre-stimulus pupil size on response velocity (BF_10_ = 6.37×10^85^) which was due to slower response velocities when pre-stimulus pupil size was small (post-hoc Bayesian t-test, BF_10_ = 1.27×10^74^ and BF_10_ = 4.98×10^27^ for small versus medium and large pupil sizes respectively). A post-hoc Bayesian t-test comparing velocity for medium and large pre-stimulus pupil sizes provided strong evidence in support of the null hypothesis that velocity did not differ (BF_10_ = 0.08). Given that velocity distributions are truncated at 0 (and are thus not normal distributions), we also conducted a classical Kruskal-Wallis Test followed by post-hoc Dunn’s tests and the results agreed with the Bayesian statistics (H=494.74, p<0.001; low versus medium: Z=-20.57, Bonferroni corrected p-value <0.001; low versus high: Z=-12.64, Bonferroni corrected p-value <0.001; medium versus high: Z=-1.70, Bonferroni corrected p-value =0.266). This pupil-vigor relationship resembles the relationship between aberrant neuronal activity and peak velocity. Therefore, increased arousal may modulate vigor by making aberrant S-R mapping activations more likely.

**Figure 4.**
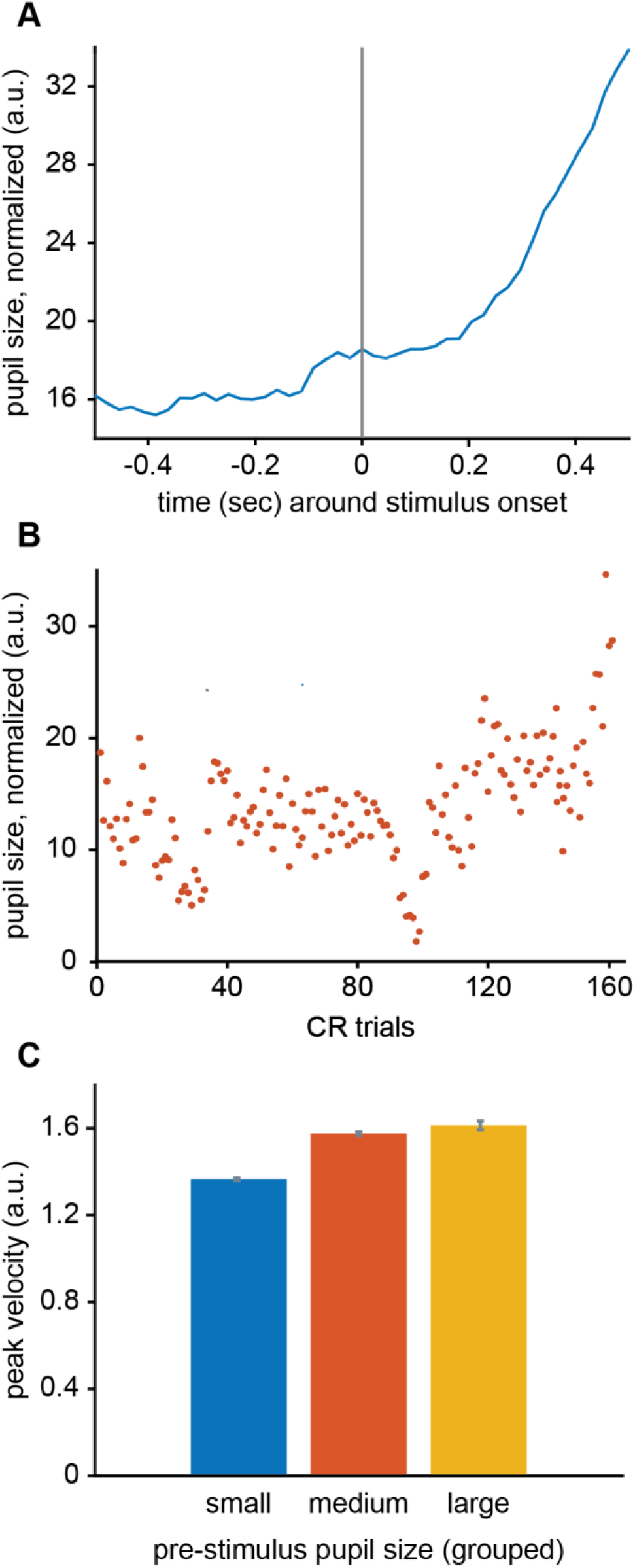
Pre-stimulus pupil size is related to peak velocity. A One example of the pupil size trace around the time of stimulus onset. **B** Averaged pre-stimulus pupil size (0.5 sec before stimulus onset until stimulus onset) across trials (one session). **C** Trials are grouped into sets with large, medium, and small pre-stimulus pupil size for all 37 rats. This was done by taking the maximum and minimum pupil size per session and subsequent splitting of this range into tertiles. The peak velocity was different for small versus medium (BF_10_ = 1.27×10^74^) and small versus large (BF_10_ = 4.98×10^27^) pre-stimulus pupil size groups. There was no evidence for a difference in peak velocity between the medium and large pupil size groups (BF_10_ = 0.08).

## Discussion

In this study, we investigated how the activity of S-R mapping encoding units in the PFC of rats relates to response initiation and response vigor. We show that when one S-R mapping is activated, in the presence of the other stimulus, the incorrect response is initiated. The vigor of this incorrect response is correlated with the strength of PFC neuronal activation. We further show that pre-stimulus arousal is related to the vigor of these incorrectly initiated responses.

Heightened arousal has been associated with committing errors and shown to modulate anterior cingulate cortex activity (Ebitz and Platt, 2015). Given that arousal-related brain regions, such as the locus coeruleus, project to the PFC in rats (and non-human primates) (Levitt et al., 1984; Chandler et al., 2014), our results suggest the possibility that arousal and neuromodulation of the PFC could promote aberrant S-R mapping activity and thus the initiation of incorrect responses (hence the association to error trials) and the vigor of those responses. It has long been postulated that arousal systems are linked to response vigor (Nieuwenhuis, 2024), but this link has not been thoroughly tested until recently (Beerendonk et al., 2024). Our results further support this link and, moreover, suggest a possible neuronal mechanism for the arousal-vigor relationship. Increased neuromodulatory system activity may bias PFC neuronal activity towards increased S-R mapping encoding activity which in turn would drive higher response vigor for the subsequent behavioral response.

The link between S-R mapping encoding and the vigor of the subsequent response is completely dependent on the learned context of this task. By analogy, humans have learned to stop in response to red traffic lights, but not all red lights. It is important to note that our findings do not suggest that PFC stimulus-evoked neuronal activity will necessitate or evoke running outside of the task context used here. In a confined context with specific response requirements, S-R mappings are an efficient representation of the task demands. Thus, S-R mappings are a building block for the variety of cases where PFC neurons become selective to task parameters, such as abstract rules (Sakagami and Tsutsui, 1999; Wallis et al., 2001; Veit and Nieder, 2013; Reinert et al., 2021), a stimulus (or stimulus feature) itself (Sakagami and Niki, 1994b; Bichot et al., 1996; Lauwereyns et al., 2001), motor preparation (Chen et al., 2017), or the value of a stimulus (Wal et al., 2021).

Our results align with previous findings in support of the Global Neuronal Workspace (GNW) theory (Dehaene et al., 1998; Mashour et al., 2020). According to this theory, conscious report of a stimulus requires that a state of “ignition” is reached in frontal cortex. This ignition state is a strong, transient increase in activity around 300 msec post stimulus. A recent study has shown that in the context of a detection task, ignition can be reached sometimes in stimulus-absent trials due to fluctuating activity in PFC (Vugt et al., 2018). On these trials the subject commits a false alarm. The PFC activity we observed also peaks around 300 msec post-stimulus and is followed by response initiation. The aberrant S-R mapping activity could thus be considered an example of the wrong “pool” of neurons reaching the state of ignition due to ongoing fluctuations of neuronal activity. Our results expand GNW theory is two new directions. First, we show that the intensity of ignition could be directly linked to the vigor of the subsequent response. Second, the fact that the response in our paradigm was aborted before the animals committed an error suggests that behaviors triggered by a state of ignition can still be stopped. This behavior most likely requires a second conscious decision to follow the first one very quickly. GNW theory predicts that this second conscious decision requires a second ignition state. In our task context, this would be akin to internally driven activation of the “correct pool” of S-R mapping selective neurons. These two expansions of the GNW theory indicate the need to investigate the interplay of different conscious decisions and hence the interplay of different states of ignition, as well as the relationship between ignition states and response vigor.

## Acknowledgements

This work was funded by the Max Planck Society, the University of Helsinki (Helsinki Institute of Life Science), the Sigrid Jusélius Foundation, and the Research Council of Finland (Project Grant, decision 358106).

